# Proteostasis is differentially modulated by inhibition of translation initiation or elongation

**DOI:** 10.1101/2022.01.23.477418

**Authors:** Khalyd J. Clay, Yongzhi Yang, Michael Petrascheck

## Abstract

Recent work has revealed an increasingly important role for mRNA translation in maintaining proteostasis. Inhibiting translation protects from various proteostatic insults, including heat, expression of aggregation-prone proteins, or aging. However, multiple studies have come to differing conclusions about the mechanisms underlying the protective effects of translation inhibition. Here, we systematically lower translation either by pharmacologically inhibiting translation initiation or elongation and show that each step activates distinct protective responses in *Caenorhabditis elegans*. Targeting initiation triggers an HSF-1 dependent mechanism that protects from heat and age-associated protein misfolding but not from proteotoxicity caused by proteasome dysfunction. Conversely, targeting elongation triggers an HSF-1 independent mechanism that protects from heat and proteasome dysfunction but not from age-associated protein aggregation. Furthermore, while inhibiting translation initiation increases lifespan in wild-type worms, inhibiting translation elongation only extends lifespan when the animals exhibit preexisting proteotoxic stress—either as a result of aggregation-prone protein expression or *hsf-1* deficiency. Together our findings suggest that organisms evolved complementary mechanisms that the mRNA translation machinery can trigger to restore proteostasis.

## Introduction

Protein synthesis is a highly regulated process involving the precise orchestration of many chaperones, co-factors, enzymes, and biomolecular building blocks. It is critical at every level of the life cycle from development through aging and is central to stress adaptation (Higuchi-Sanabria et al., 2018). Protein synthesis largely determines the folding load on the proteostasis network, which regulates protein production, folding, trafficking, and degradation to maintain a functional proteome. The imbalance of the proteostasis network caused by age-associated stress or acute environmental insults leads to misfolding of proteins and the accumulation of aggregates, and eventually to disease (Balch et al., 2008).

A substantial body of work has revealed that the protein synthesis machinery directly participates in protein folding or aggregation. For example, in mice, point mutations in specific tRNAs or components of the ribosomal quality control (RQC) pathway can lead to protein aggregation and neurodegeneration (Chu et al., 2009; Nedialkova and Leidel, 2015; Nollen et al., 2004; Vo et al., 2018; Yonashiro et al., 2016). Similarly, early RNAi screens in the nematode *C. elegans* identified several ribosomal subunits whose knockdown increased aggregation of polyglutamine (polyQ) proteins (Nollen et al., 2004). These findings highlight the importance of translation in protein misfolding.

However, subsequent studies reveal a more intricate role of translation in protein aggregation. Depending on how translation is modulated can lead to either decreased or increased protein aggregation. For example, RNAi-mediated knockdown of translation initiation factors increase lifespan, improve proteostasis, and reduce protein aggregation in *C. elegans* (Balch et al., 2008; Lan et al., 2019; McQuary et al., 2016; Rogers et al., 2011). Similarly, work in yeast and cell culture shows that pharmacological inhibition of translation prior to a heat shock prevents proteins from aggregating (Choe et al., 2016; Medicherla and Goldberg, 2008; Riback et al., 2017; Xu et al., 2016). While studies in *C. elegans*, cell culture, and yeast agree that inhibition of translation reduces protein aggregation, their proposed underlying mechanisms differ. Overall, the proposed mechanisms can be categorized into two broad models on how lowering translation reduces protein aggregation.

The first model, referred to as the *selective translation model*, proposes that inhibition of translation is selective. In the selective translation model, the increased availability of ribosomes leads to differential translation of mRNAs coding for stress response factors and thus to increased folding capacity (Lan et al., 2019; McQuary et al., 2016; Rogers et al., 2011; Seo et al., 2013). In general, studies proposing a version of *the selective translation* model show that inhibiting translation requires HSF-1 to reduce protein aggregation (Tye and Churchman, 2021). In the *selective translation model*, protein aggregation is reduced by an HSF-1 dependent active generation of folding capacity to remodel the proteome.

The second model, referred to as the *reduced folding load model*, proposes that newly synthesized proteins are the primary aggregation–prone species of proteins. Therefore, newly synthesized proteins constitute the most significant folding load on the proteostasis machinery. Inhibition of translation reduces the concentration of newly synthesized proteins and thus the load on the folding machinery. In contrast to the *selective translation model*, which proposes selective protein synthesis of HSF-1 dependent stress response factors, the *reduced folding load model* does not depend on specific factors but generates folding capacity by reducing the overall folding load. A fundamental problem comparing previous studies has been their use of different model organisms, different proteostatic insults, and modes of inhibition, each of which could potentially account for the varied conclusions.

In this study, we systematically compared these two models in *C. elegans* using pharmacological agents to block various steps along the protein production cycle. We characterize how lowering translation protects *C. elegans* from proteotoxic insults such as proteasome inactivation, heat shock, and aging. We find evidence for both the *selective translation* and the *reduced folding load models*. Our data reveal that the step inhibited in mRNA translation dictates which of the two protective mechanisms is activated. Furthermore, we show that the two mechanisms are complementary in protecting from proteostatic insults suggesting that a cell may activate one over the other depending on the proteostatic stress. Overall, we provide evidence that elongation inhibitors ameliorate ongoing toxicity by *reducing folding load*. In contrast, initiation inhibitors elicit an active mechanism that prevents damage by remodeling the proteome dependent on HSF-1 and a likely increase in protein turnover via the proteasome.

## Results

### Translation inhibitors improve thermotolerance

We first set out to identify suitable pharmacological initiation and elongation inhibitors by screening a series of translation inhibitors (Table 1) for their ability to inhibit protein synthesis in *C. elegans* (Dmitriev et al., 2020). To monitor protein synthesis, we employed SUrface SEnsing of Translation (SUnSET) (Arnold et al., 2014). In this method, translating ribosomes incorporate puromycin into newly synthesized proteins. The level of puromycin incorporation serves as a quantitative measure for translation and is detected by Western Blotting using an anti-puromycin monoclonal antibody. All six molecules (Table 1) reduced puromycin incorporation relative to DMSO controls to varying degrees (Figure 1A & B). Based on the similar inhibitory effects, we selected the two initiation inhibitors—4E1RCat and salubrinal—and the two elongation inhibitors—anisomycin and lycorine—to investigate how chemically targeting two steps of the translation cycle will affect stress–induced protein aggregation.

**Table 1:**
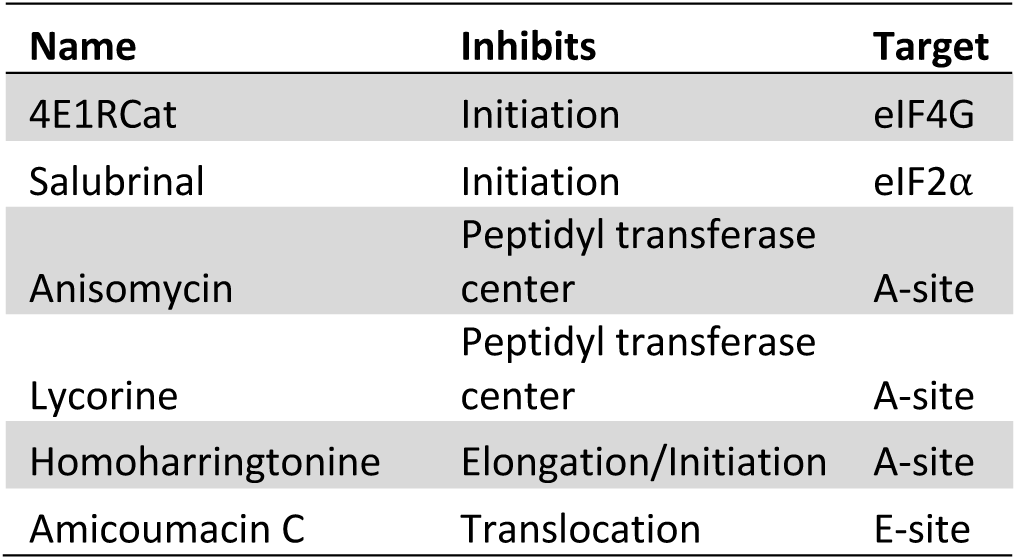
Translation inhibitors and their effects.

| Name | Inhibits | Target |
| --- | --- | --- |
| 4E1RCat | Initiation | eIF4G |
| Salubrinal | Initiation | eIF2α |
| Anisomycin | Peptidyl transferase center | A-site |
| Lycorine | Peptidyl transferase center | A-site |
| Homoharringtonine | Elongation/Initiation | A-site |
| Amicoumacin C | Translocation | E-site |

**Figure 1:**
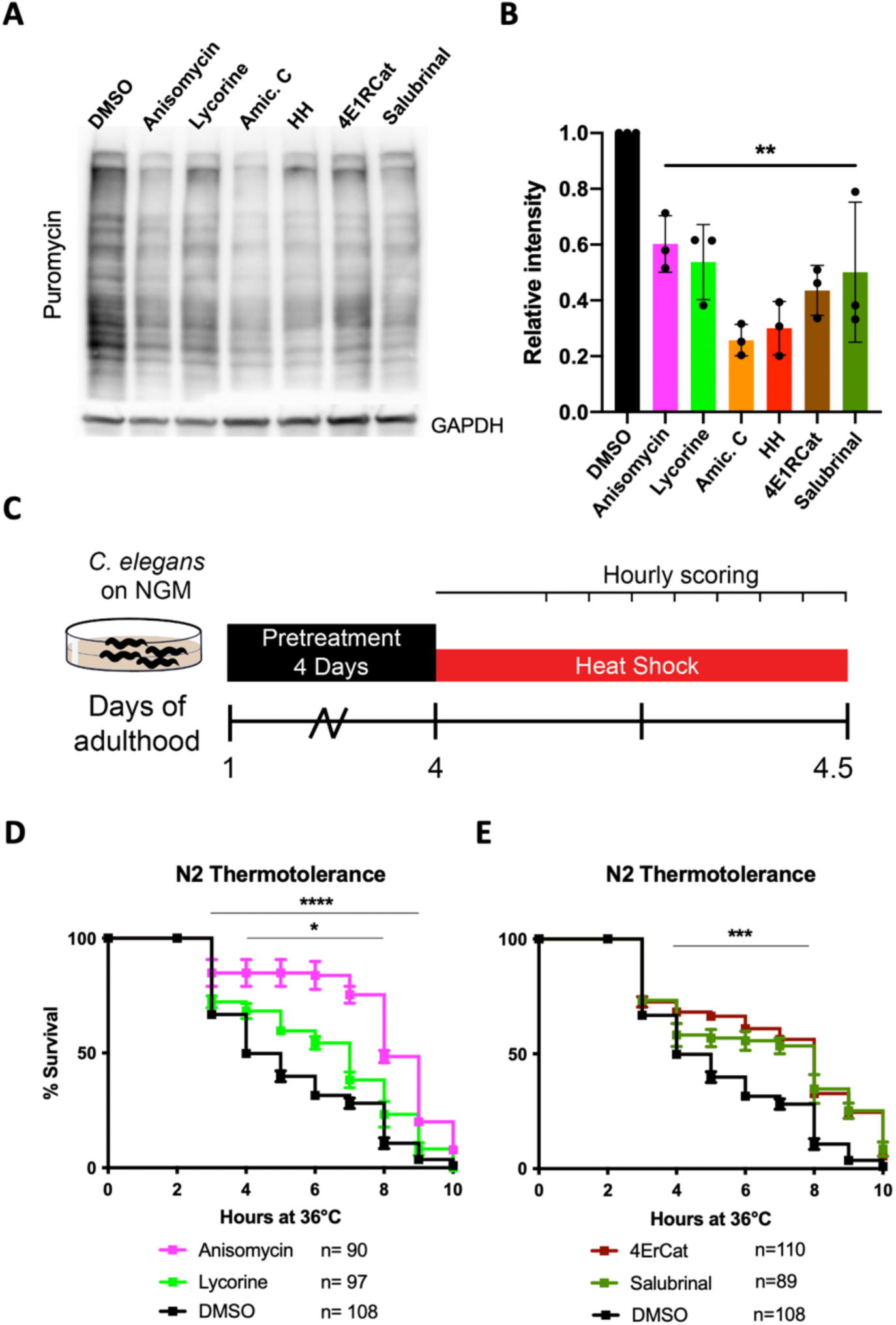
Translation inhibitors improve thermotolerance. **A)** Monitoring changes in protein synthesis using the SUnSET method. *C. elegans* were treated with solvent (DMSO) or the indicated inhibitors (100 μM) for 12 hours, followed by a 4-hour puromycin incorporation. Worms were lysed, and protein extracts were run on SDS-PAGE gels, followed by staining with puromycin antiserum. GAPDH was used as a loading control. **B)** Quantification of three independent SUnSET experiments. Significance was determined by one-way ANOVA with Dunnett’s multiple comparisons test where ** = p ≤ 0.01 Error bars indicate mean ± SD from three independent trials. **C)** Day 1 adult wild-type (N2) animals were treated for 4 days, then transferred to NGM plates. Animals were then subjected to a constant, non-permissive temperature of 36°C and scored alive/dead every hour by movement following a gentle tap. **D)** Graph shows survival as a function of hours at 36 °C of day 4 adult wild-type (N2) animals. The animals were pretreated with either DMSO (solvent control), anisomycin or lycorine. Data are displayed as mean ± SEM from three independent trials and * = p ≤ 0.05 and **** = p ≤ 0.0001 by row–matched two–way ANOVA with Šídák multiple comparisons test. **E)** Same as in Figure 1D but showing initiation inhibitors 4E1RCat and salubrinal. Data are displayed as mean ± SEM from three independent trials and *** = p≤ 0.0002 by row–matched two–way ANOVA with Šídák multiple comparisons test.

We chose thermal stress as the first stressor to induce protein aggregation and asked if both initiation and elongation inhibitors improve stress resistance. The animals were treated with either of the four inhibitors on day 1 of adulthood, and after 72 hours, treatment moved to a non–permissive temperature of 36 °C (Figure 1C). Hourly monitoring revealed that all four molecules significantly improved the survival of N2 animals (Figure 1D & E). These data show that both the inhibition of translation initiation and elongation protect from thermal stress.

### Initiation and elongation inhibitors protect from thermal stress by HSF-1 dependent and independent mechanisms, respectively

We next asked if translation inhibitors require the canonical heat shock response (HSR) controlled by the transcription factor HSF-1 to protect from thermal stress. Previous work by us and others resulted in contradictory findings on whether protection from heat by translation inhibition depends on HSF-1 (Seo et al., 2013; Solis et al., 2018; Zhou et al., 2014). Therefore, to test if translation inhibition protects from thermal stress in an HSF-1 dependent or independent manner, we repeated the thermotolerance assay in HSR–deficient *hsf-1(sy441)* mutants (Figure 2A). Only the elongation inhibitors anisomycin and lycorine protected *hsf-1(sy441)* from HS–induced proteotoxicity (Figure 2B). The initiation inhibitors 4E1RCat and salubrinal did not (Figure 2C). These results were surprising as they showed that different modes of translational inhibition activate distinct protective proteostasis mechanisms. However, this also reconciles previous contradictions as different groups inhibited translations using inhibitors with specificity for either step.

**Figure 2:**
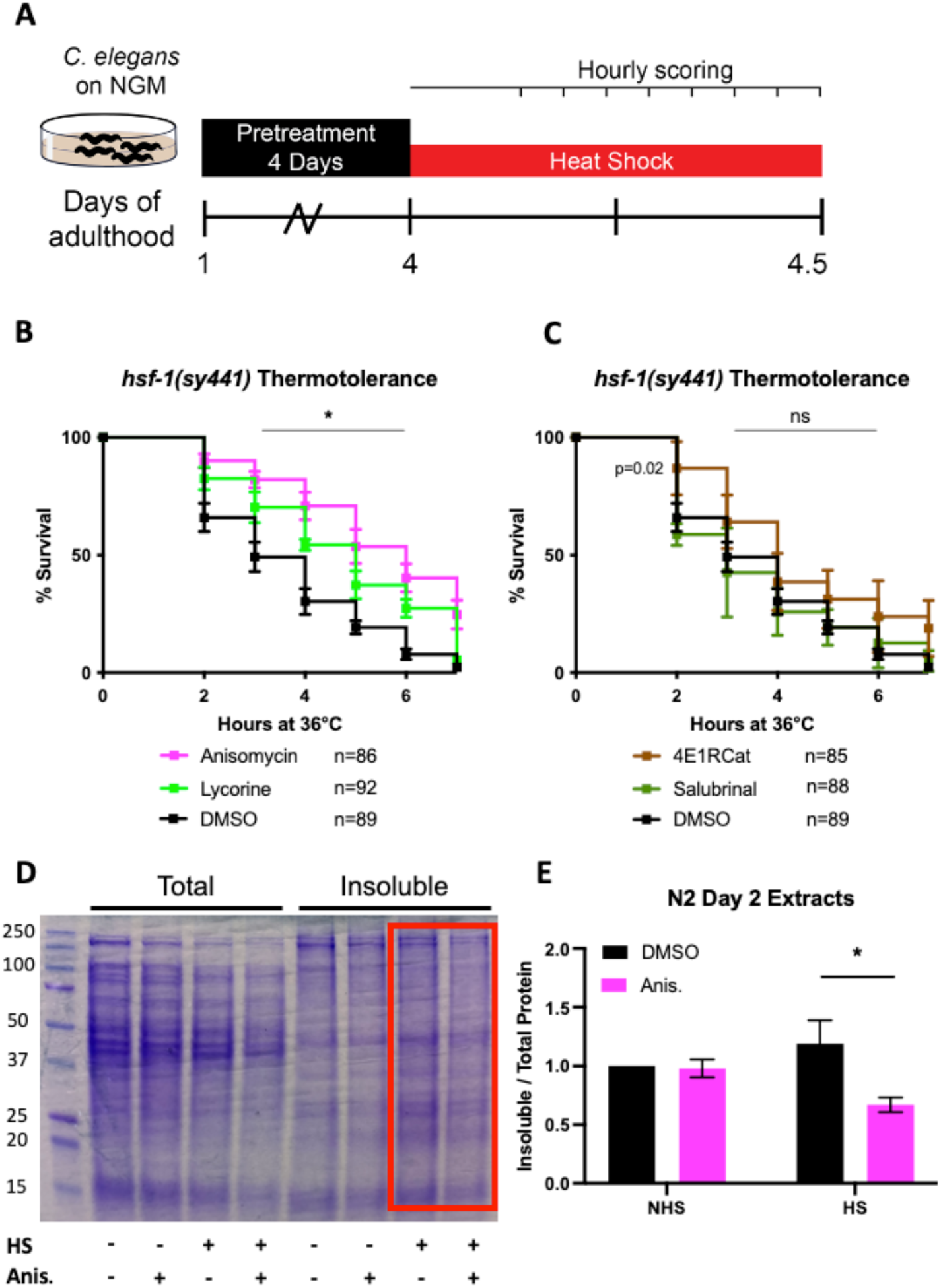
Initiation but not elongation inhibitors depend on HSF-1 to protect from thermal stress. **A)** Day 1 adult *hsf-1(sy441)* animals were treated for 4 days, then transferred to NGM plates. They were then subjected to a constant, non-permissive temperature of 36°C and scored alive/dead every hour by movement following a gentle tap. **B)** Thermotolerance assay: Graph shows survival as a function of hours at 36 °C of day 4 adult *hsf-1(sy441)* animals pretreated with either anisomycin or lycorine. Data show the mean ± SEM from three independent trials and * = p≤ 0.05 by row–matched two–way ANOVA with Šídák multiple comparisons test. **C)** Same as in Figure 2B but showing initiation inhibitors 4E1RCat and salubrinal. Data are displayed as mean ± SEM from three independent trials and * = p≤ 0.05 by row–matched two–way ANOVA with Šídák multiple comparisons test. **D)** Representative SDS-PAGE gel stained with the protein stain Comassie Blue for visualization. Anisomycin (Anis.) reduces the proportion of detergent-insoluble protein following a 2 hour HS of N2 animals. Proteins were detergent extracted, ultracentrifuged, and the insoluble pellet was resuspended in 8M urea before running on the gel. **E)** Quantification of 4 separate extractions shows anisomycin significantly reduces HS–induced aggregation in wild-type N2 animals. Gels were stain with Sypro Ruby. Data are displayed as mean ± SEM and * = p≤0.05 by two-tailed students t-test.

To confirm the ability of elongation inhibitors to protect from thermal proteotoxicity through improved proteostasis broadly, we conducted sequential detergent extractions to biochemically isolate and quantify soluble and insoluble proteins in wild-type (N2) animals (David et al., 2010; Reis-Rodrigues et al., 2012; Simonsen et al., 2008). Following a 2 hour HS at 36 °C, we observed a substantial increase in insoluble protein compared to non-heat shocked controls. Furthermore, pre-treatment with the elongation inhibitor anisomycin suppressed the increase in protein insolubility (Figure 2D and E) consistent with the observed thermal protection. Using the *hsp-16*.*2*::GFP reporter strain, we further confirmed that anisomycin treatment does not activate the HSR, either alone or in combination with a heat shock (see below, Fig. 4E). These results reveal that translation initiation inhibitors trigger an HSR dependent mechanism to protect from thermal stress, while translation elongation inhibitors trigger an HSR independent mechanism. We concluded that different modes of translational inhibition protect the proteome by genetically distinct mechanisms.

### Translational elongation, but not initiation inhibitors protect from proteasome dysfunction

The data thus far suggests that inhibition of translation protects from thermal stresses by at least two mechanisms, one that involves HSF-1 and one that does not. We, therefore, investigated whether the protective mechanisms can be further distinguished by their capacity to protect from different proteostatic insults. One main proteostasis mechanism by which cells clear protein aggregates is the proteasomal system. The proteasomal system ubiquitinylates misfolded proteins by ubiquitin ligases to target them for degradation by the 26S proteasome. Blocking proteasome degradation by bortezomib, a specific inhibitor of the 20S subunit, results in the formation of protein aggregates and proteotoxic stress (Schrader et al., 2016).

Proteasomal stress induced by the treatment with bortezomib killed ~40% of the wild-type N2 animals by day 8. Cotreatment of bortezomib with any of the two translation initiation inhibitors enhanced proteasome toxicity leading to over 80% of the animals dying, while cotreatment with elongation inhibitors improved morphology but had no effect on survival (Figure 3B). Thus, while initiation inhibitors protect from thermal stress, they sensitize the animals to proteasomal stress.

**Figure 3:**
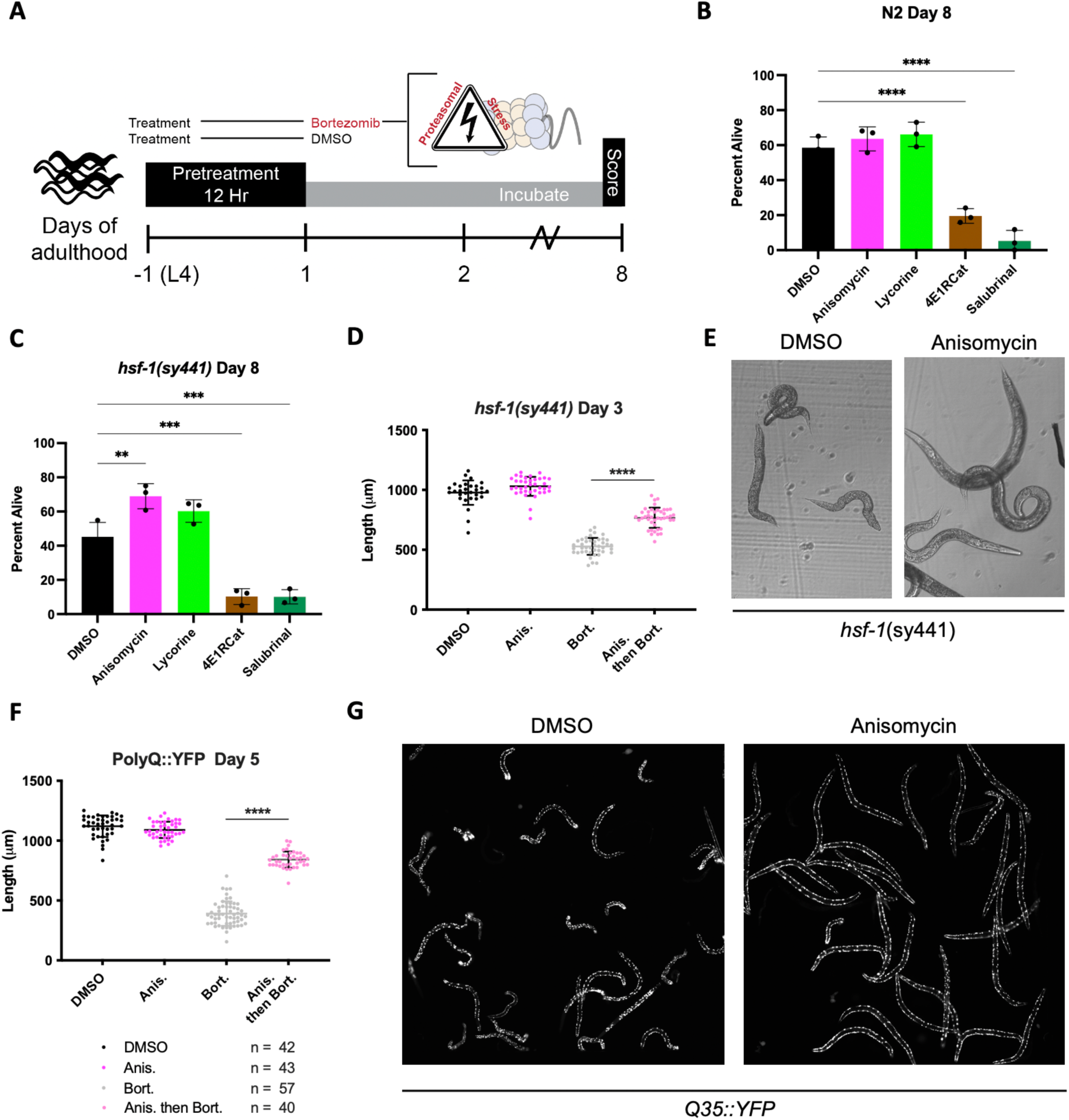
Elongation inhibitors protect from proteasomal stress independent of *hsf-1*. **A)** Experimental strategy for treating animals with translation inhibitors followed by inhibition of the 20S proteasome by bortezomib. Worms are pretreated for 12 hours with DMSO or indicated inhibitors, followed by bortezomib (75 μM) treatment. The animals were then incubated with the combined treatment for 8 days and scored as alive/dead based on movement. **B)** Elongation inhibitor treatment improved morphological phenotypes (not shown) but did not protect from bortezomib-induced proteotoxicity in N2 animals. Initiation inhibitor treatment enhanced toxicity of proteasomal stress. Data are displayed as mean ± SD and **** = p< 0.0001 by one-way ANOVA. **C)** In *hsf-1(sy441)* animals, elongation inhibitors protected from bortezomib-induced proteotoxicity while initiation inhibitors continue to sensitize worms to the resulting proteotoxicity. Data are displayed as mean ± SD and ** = p<0.01 by one-way ANOVA. **C)** Measured length of *hsf-1(sy441)* worms at day 3 of adulthood. Anisomycin treatment almost completely rescued the *sma* phenotype induced by bortezomib. Data are displayed as mean ± SD and **** = p<0.0001 by a two-tailed unpaired t-test. **D)** Representative brightfield images of day 3 *hsf-1(sy441)* animals treated with the indicated compounds as outlined in figure 5A. Anisomycin pre-treatment prevented the *sma* phenotype observed to be caused by proteasomal inhibition. **E)** Measured length of PolyQ worms at day 5 of adulthood. Anisomycin treatment almost entirely rescued the small (*sma*) phenotype induced by bortezomib. Data are displayed as mean ± SD and **** = p<0.0001 by a two-tailed unpaired t-test. **F)** Representative fluorescent images PolyQ worms at day 5 of adulthood treated with the indicated compounds as outlined in Figure 5A. Bortezomib treatment caused the worms to contract into a *sma* and *unc* phenotype (left panel), and anisomycin pre-treatment prevented these pathological phenotypes (right panel).

Next, we repeated the bortezomib-induced proteasomal stress survival experiments using strains with reduced protein folding capacity—either because of a mutation in *hsf-1* or because expression of an aggregation-prone protein (PolyQ35::YFP). Treatment of *hsf-1(sy441)* mutant animals with bortezomib killed ~60% of the *hsf-1(sy441)* animals, a substantial increase in toxicity compared to N2. Cotreatment with translation initiation inhibitors again enhanced proteasome toxicity leading to over 80% of the animals to die. Cotreatment with elongation inhibitors, however, suppressed bortezomib toxicity and enabled about 70% of the *hsf-1(sy441)* animals to survive (Figure 3C). This survival rate was close to what was seen in bortezomib–treated wild-type animals, suggesting that elongation inhibitors mostly rescue the *hsf-1(sy441)* proteostasis phenotype. We also observed bortezomib-treated *hsf-1(sy441)* animals to shrink in size (*sma* phenotype) and to become severely uncoordinated (*unc* phenotype). Treatment with anisomycin substantially rescued *sma* and *unc* phenotypes, almost completely reversing the animals back to normal size (Figure 3D & F).

The AM140 strain expresses a stretch of 35 glutamine residues fused to YFP (PolyQ::YFP) in the muscle, which we used as a second model of reduced protein folding capacity. Expression of the aggregation-prone PolyQ stretch increases the protein folding load on the proteostasis system. Thus, its aggregation propensity acts as a sensor for protein folding capacity (Brignull et al., 2006; Moronetti Mazzeo et al., 2012). As expected, treatment of PolyQ::YFP animals with bortezomib caused extensive protein aggregation, a reduction in body size (*sma* phenotype), and an uncoordinated phenotype (*unc* phenotype). In addition, inhibition of translation initiation again exacerbated bortezomib toxicity but was not further quantified. In contrast, inhibition of elongation by anisomycin almost entirely rescued both the *sma* and *unc* phenotypes revealing protection from proteasome toxicity in the context of reduced folding capacity (Figure 3F &G).

We concluded that inhibition of translation initiation exacerbates bortezomib-induced proteasome stress, probably because its downstream protective mechanism heavily depends on proteasomal degradation (see discussion). We further concluded that elongation inhibition provided only limited direct protection from proteasomal stress since we did not see an increase in survival in wild-type animals. However, elongation inhibitors were highly protective from bortezomib toxicity in strains with reduced protein folding capacity, suggesting that elongation inhibitors protect by reducing the need for protein folding capacity by lowering the concentration of newly synthesized proteins.

### Translational elongation and initiation inhibitors protect from protein aggregation

The ability of anisomycin to mitigate bortezomib toxicity in PolyQ::YFP animals suggested that translation elongation inhibitors free up folding capacity through the reduction of overall protein synthesis. To monitor folding capacity *in live imaging*, we developed a live imaging experimental procedure to examine how translation inhibition will dynamically affect protein aggregation in real-time (Figure 4A). After a 2 hour HS, the initially diffuse PolyQ signal gradually localized into puncta (Figure 4B, Video 1). Aggregation foci formation resulted from a redistribution of the YFP signal into aggregation foci as the total level of YFP fluorescent signal for a given animal remained constant (S1).

**Figure 4:**
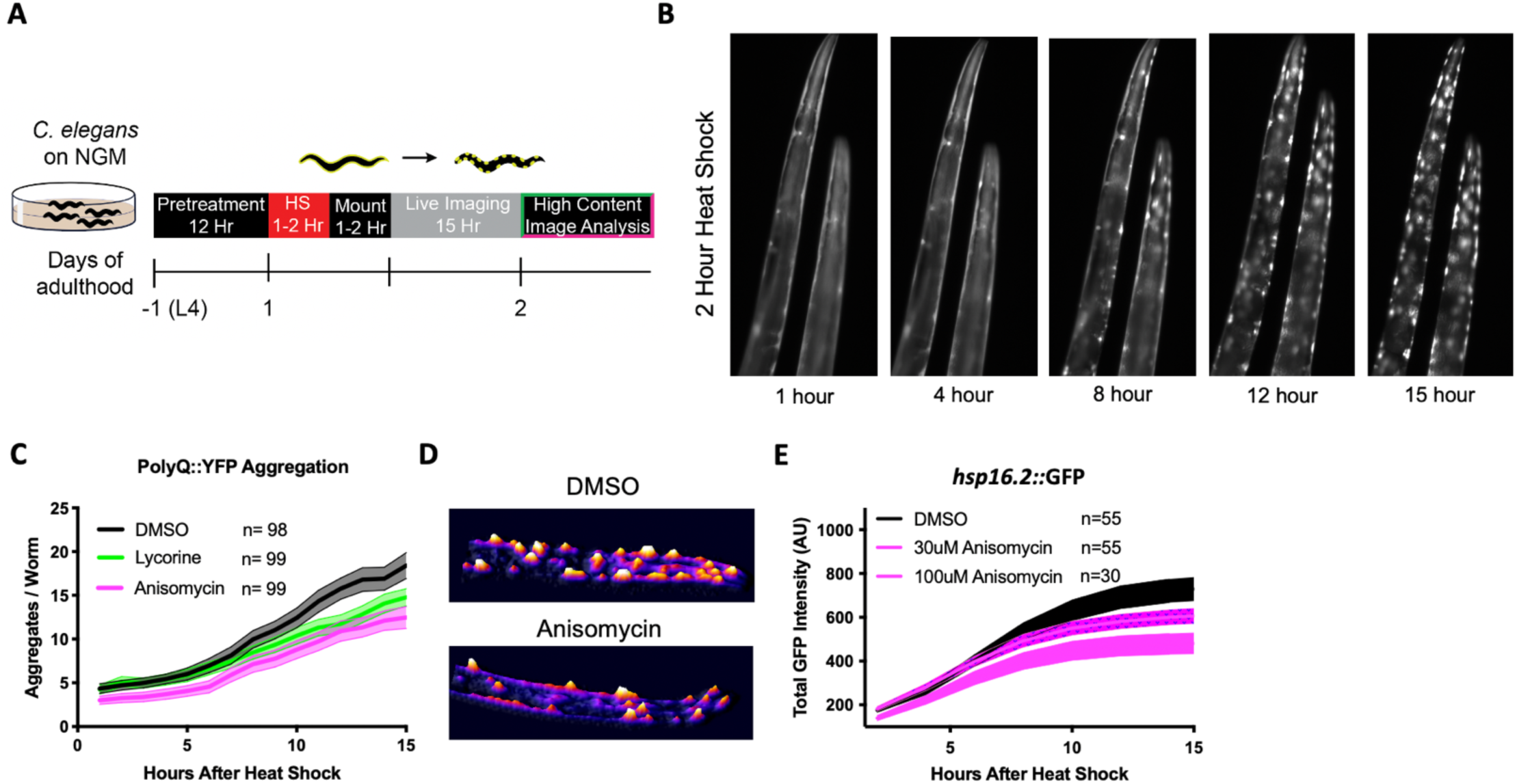
Elongation inhibition protects from heat shock-induced protein. **A)** Day 1 AM140 adult worms expressing the polyglutamine-YFP fusion protein (PolyQ::YFP) in their muscle were subjected to HS on NGM plates for 2 hours at 36 °C followed by a 1-2 hour mounting/immobilization procedure in 384 well plates and subsequent live imaging for 15 hours. **B)** Fluorescent time-lapse images of two animals expressing the PolyQ::YFP fusion protein in their muscle. The animals were embedded in the hydrogel for immobilization. Following a 2-hour HS, animals were imaged over 15 hours; by 8 hours, the YFP signal began to localize into discrete puncta that persisted through the observation time. Overall fluorescence remained stable (see S2). **C)** Graph shows the mean number of PolyQ aggregates per worm as a function of time following HS. *C. elegans* (PolyQ::YFP) were pretreated with lycorine, anisomycin (100 μM), or DMSO. Lines indicate mean, and shading indicates 95% CI. **D)** (Top) Representative images of control and 100 μM anisomycin treated PolyQ animals 15 hours after HS (Bottom). The representative images shown have been uniformly modified using the ‘3D Surface Plot’ plugin in ImageJ to visualize aggregates. **E)** Dose-dependent suppression of HS-induced *hsp-16*.*2*::GFP reporter activation (Day 1 animals) after 12 hour pretreatment of anisomycin. 95% CI, as indicated by shading.

Only elongation inhibitors lycorine and anisomycin significantly reduced HS–induced polyQ aggregation, while the initiation inhibitors 4E1RCat and salubrinal did not (Figure 4C and S2). Pre-treatment with anisomycin and lycorine reduced the number of polyQ aggregates per worm. However, pre-treatment did not change the onset of aggregation, the rate of formation, or the final size of the aggregation foci (Morley et al., 2002) (Figure 4D). Furthermore, we found the effect of anisomycin or lycorine to be time-dependent. They strongly inhibited aggregation following a 12 hour preincubation period but less so following a 4-hour preincubation (S3). Starting treatment with anisomycin after the HS did not reduce the number of aggregation foci (not shown). These results suggest that anisomycin and lycorine reduce the early formation of aggregate foci but do not alter the dynamics once aggregation begins. Anisomycin dose-dependently decreased the activation of the HSR, as measured by the *hsp-16*.*2*::GFP reporter strain, confirming again that protection from aggregation is independent of *hsf-1* (Figure 4E, S4).

### Lifespan extension by translational elongation and initiation inhibitors is dictated by genetic background

Lowering translation is an established mechanism to extend lifespan and delay aging (Anisimova et al., 2018; Klaips et al., 2018; Steffen and Dillin, 2016). Furthermore, aging is a well-known driver of protein aggregation. However, to our knowledge, it has never been investigated if the anti-aggregation and anti-aging effects of translational inhibition can be uncoupled and if the mode of translational inhibition influences these phenotypes. Inhibition of translation in wild-type animals, using the two elongation inhibitors anisomycin and lycorine, showed no, or only a minor lifespan extension in N2 animals (Fig 5A). In contrast, inhibition of translation by the initiation inhibitors 4E1RCat and salubrinal dose-dependently extended lifespan (Fig. 5B). This difference was observed despite that all four translation inhibitors reduced protein translation to the same extent (Fig. 1A). Thus, the difference in the effect on lifespan by elongation inhibitors and initiation inhibitors cannot be explained by reducing overall protein synthesis alone.

**Figure 5:**
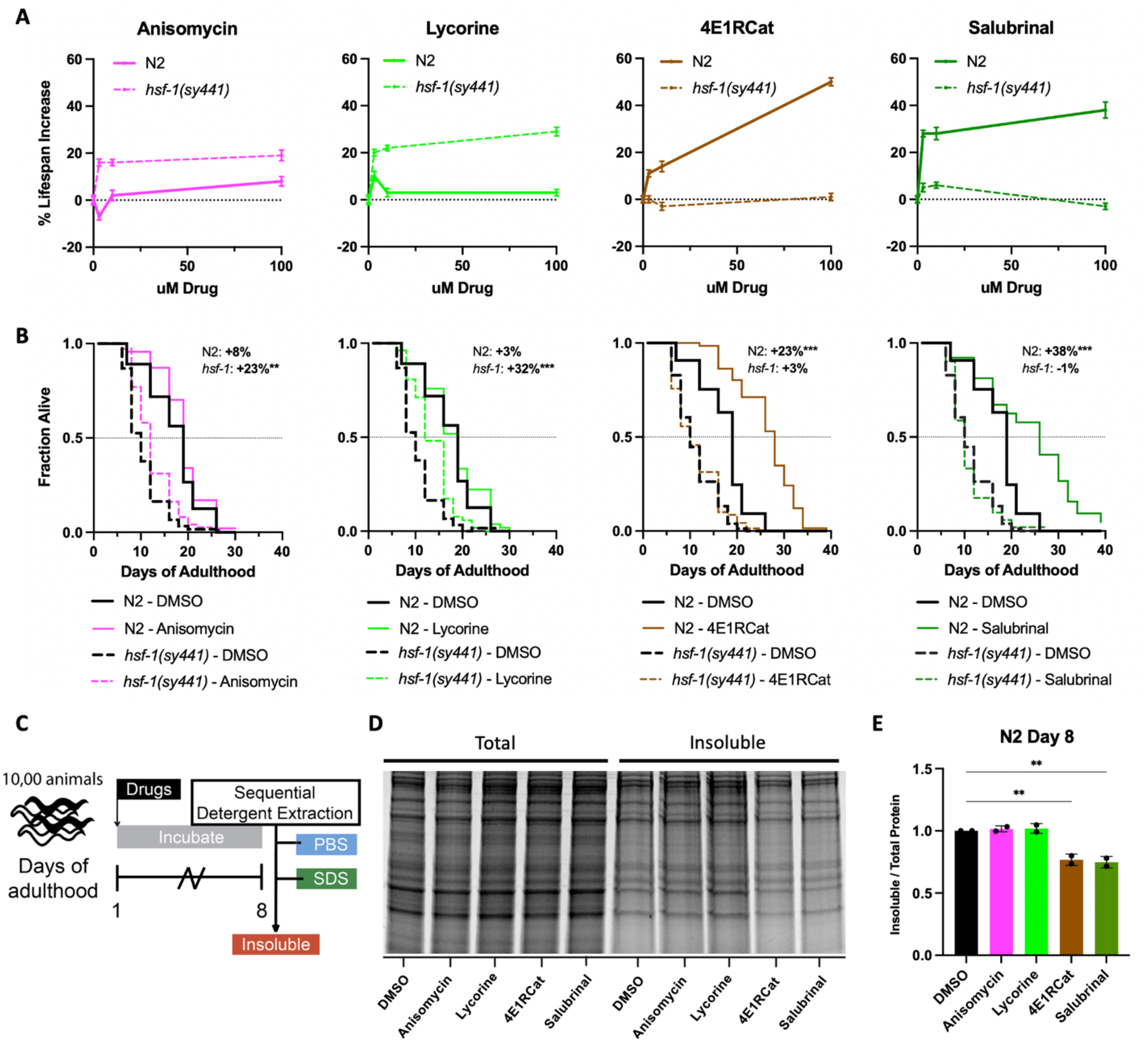
Reciprocal lifespan extension by translation inhibitors in *hsf-1(sy441)* animals. **A)** Mean lifespan as a function of translation inhibitor concentration. A maximum increase in lifespan of N2 animals (dashed black line) was seen at a 100 μM concentration. The lifespan of *hsf-1(sy441)* was tested in parallel, where the effects of initiation versus elongation inhibitors were reversed. Error bars indicate ± SEM. See Supplementary Table 1 for details. **B)** Survival curves from representative experiments show the fraction of wild-type (N2, solid line) or *hsf-1*–deficient (dashed line) animals when treated with 100 μM of the indicated compound. Black lines indicate DMSO treatment, and colored lines indicate inhibitor treatment. Data are displayed as a Kaplan-Meier survival curve and significance determined by log-rank test. **C)** Experimental strategy for treating animals with inhibitors and isolating detergent-insoluble fractions. First, 10,000 animals were treated and allowed to age for 8 days before being washed with M9, frozen in N_2_, and mechanically lysed. Then proteins were extracted from the total lysate based on solubility, and an aliquot from each fraction was run on an SDS-PAGE gel. **D)** Representative SDS-PAGE gel stained with the protein stain Sypro Ruby. 4E1RCat and salubrinal reduce the amount of age-associated protein aggregation. **E)** Quantification of 2 separate experiments shows 4E1RCAt and salubrinal to significantly reduce age-associated aggregation in wild-type (N2) animals. Data are displayed as mean ± SEM and ** = p < 0.01 by two-tailed students t-test.

**Figure 6:**
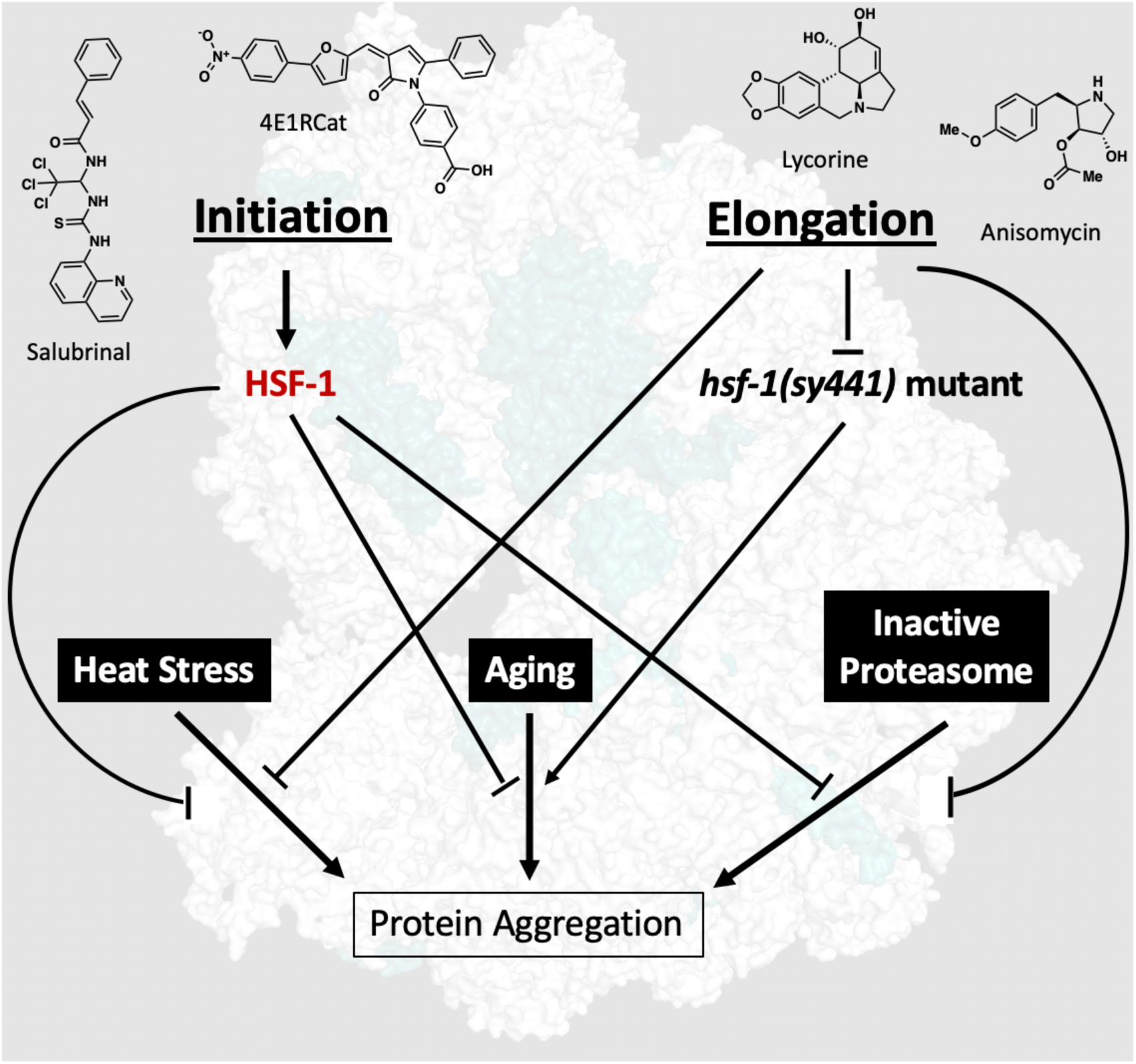
Consequences of proteotoxicity on aggregation at the ribosomal interface. Initiation inhibitors, salubrinal, and 4E1RCat require *hsf-1* to elicit a protective response following proteotoxic stress, functioning similar, if not overlapping with, canonical stress responses. Conversely, elongation inhibitors, anisomycin, and lycorine do not boost protection in healthy wild–type animals but rescue proteotoxicity in animals with preexisting proteostasis deficits such as *hsf-1* mutants or animals expression misfolded proteins.

As the ability of the initiation inhibitors 4E1RCat and salubrinal to protect from heat stress was dependent on the transcription factor HSF-1, we asked if they were able to extend the lifespan of *hsf-1(sy441)* mutants. Both 4E1RCat and salubrinal failed to significantly extend lifespan in *hsf-1(sy441)* mutants. This result is consistent with the model put forward by Rodgers et al., which proposes that inhibition of translation initiation extends lifespan by lowering overall translation while selectively promoting the translation of cap-independent transcripts, many of which are transcribed by HSF-1.

However, to our surprise, treatment with the elongation inhibitors anisomycin and lycorine extended the lifespan of *hsf-1(sy441*) mutants by ~20% (Fig. 5B). We interpret this lifespan extension as a partial rescue of the protein folding defect in *hsf-1(sy441*) mutants as the increase in lifespan did not reach the lifespan of wild-type animals. Taken together, however, our data demonstrate that longevity induced by translation inhibition subsumes several different mechanisms that lead to longevity.

We next tested the ability of all four inhibitors to reduce age-associated protein aggregation. If inhibition of translation alone is sufficient to reduce protein aggregation independently of any downstream mechanisms, all four inhibitors should reduce age-associated protein aggregation. Conversely, if the reduction of age-associated protein aggregation is closely linked to longevity, then only the initiation inhibitors should reduce protein aggregation. We treated day 1, N2 animals with 100 μM of each of the four inhibitors and allowed the animals to age for eight days, after which we separated proteins based on solubility (Figure 5C). We found that only the two initiation inhibitors that extended lifespan caused significant decreases in the amount of SDS–insoluble aggregates and that the elongation inhibitors failed to do so (Figure 5D & E).

Taken together, these data support a model in which inhibition of translation elongation rescues proteostasis–compromised animals by reducing folding load but without generating additional protein folding capacity. Separately, inhibition of translation initiation protects and improves longevity through selective translation that activates an HSF-1 dependent mechanism. However, inhibition of translation initiation does not rescue or even can further exacerbate damage in animals with preexisting proteostasis imbalance.

## Discussion

In this study, we set out to answer how translation inhibition exerts its beneficial effects on proteostasis and longevity in the metazoan *C. elegans*. We considered two previously proposed models (Medicherla and Goldberg, 2008; Rogers et al., 2011; Seo et al., 2013; Zhou et al., 2014). The *selective translation model* proposes that inhibition of translation improves proteostasis by an active mechanism remodeling the proteome through differential translation. In the selective translation model, the system increases folding capacity through the increased translation of HSF-1 targets and by proteasomal degradation of preexisting proteins. The second model, the *reduced folding load model*, proposes that inhibition of translation improves proteostasis by lowering the concentration of newly synthesized proteins, thereby increasing relative folding capacity via a reduced folding load on the proteostasis machinery.

Our data establish evidence for both models. We show that the mode of translation inhibition determines the mechanism of proteome protection. Translation initiation inhibitors initiate a selective translation mechanism that requires HSF-1 and the proteasome (Howard et al., 2016; Rogers et al., 2011; Seo et al., 2013). In contrast, elongation inhibitors reduce folding load and do not require HSF-1 or the proteasome. Elongation inhibitors rescue deficiencies induced by the lack of these components. What was particularly striking to us was how cleanly different modes of translational inhibition could be separated based on the stressor. The most general conclusion that we draw from our study is that inhibition of translation can trigger genetically and biochemically distinct, HSF-1 dependent and independent proteostasis mechanisms.

Translation can be separated into the three major steps, initiation, elongation, and termination, each of which can be further subdivided into several minor steps. Translation is further regulated by signaling factors and ribosome assembly rates, which are also subject to proteostatic intervention. Our data show that the phenotypic consequences of lowering translation depend on the environmental context and the inhibited step of translation. It remains to be seen if modulation of additional steps, such as termination, triggers additional genetically distinct proteostasis mechanisms.

Inhibiting either initiation or elongation protects from the damaging effects of heat shock. Otherwise, the two protective mechanisms are complimentary, with one being protective while the other is not. Overall, our data suggest that initiation inhibitors remodel the proteome in an HSF-1 and proteasome-dependent manner, as previously suggested. In contrast, elongation inhibitors act independently of both HSF-1 and the proteasome, thereby rescuing the phenotypic consequences of their inactivation. The most striking example of this dichotomy was seen when preexisting folding problems—caused by the lack of either HSF-1 or the expression of the aggregation-prone polyQ protein—were exacerbated by the chemical inhibition of the proteasome. These combined insults led to increased protein aggregation, uncoordination, shrinking body size, and death of the animals. Inhibition of elongation almost entirely rescued these phenotypes. In sharp contrast, inhibition of initiation made the animals significantly worse and increased death. Our data reveal that the mode by which translation is inhibited leads to the activation of distinct downstream mechanisms that protect the organism from different insults.

A second striking example observed was the ability of initiation inhibitors to extend lifespan while the elongation inhibitors did not. A long-standing question in the field of longevity is if lowering protein synthesis by inhibiting translation is sufficient to extend lifespan. Alternative explanations for the observed longevity include lowering the expression of specific proteins (e.g., DAF-2) or the activation of a specific downstream mechanism that is activated by low translation rates (Howard et al., 2016). Our data suggest some proteome remodeling may be necessary to extend lifespan (Koyuncu et al., 2021).

The lack of longevity from anisomycin and lycorine treatment in our experiments is unlikely to be the result of toxicity due to an off-target effect. First, lycorine and anisomycin are structurally very distinct and are unlikely to share off-targets. Second, and more importantly, both extend the lifespan of *hsf-1(sy441)* mutants, which strongly argues against a toxic side effect. In general, we observed that elongation inhibitors are good at rescuing defects in strains with impaired proteostasis. In strains with impaired proteostasis, inhibition of elongation lowers the production of newly synthesized proteins and thus reduces the folding load on the proteostasis machinery. This effect alleviates folding problems and thus allows the elongation inhibitors to rescue the shortened lifespan of proteostasis–deficient strains.

### Ideas and Speculation

The ability of elongation inhibitors to improve proteostasis in strains with preexisting folding defects on the organismal level raises the interesting question of whether they could be developed therapeutically. It has often been suggested that the protein degradation machinery such as autophagy or the proteasome is overloaded in protein folding diseases. If the *C. elegans* results translate to mammals, treating a mouse model of a protein-folding disease with an initiation inhibitor would be detrimental. In contrast, treatment with an elongation inhibitor would be beneficial. Furthermore, these *C. elegans* findings may explain some surprising previous observations. For example, treating the SOD1 G93A mouse model of amyotrophic lateral sclerosis (ALS) with rapamycin, a drug modulating translational initiation through the phosphorylation of 4E-BP1 exacerbates ALS disease phenotypes (Zhang et al., 2011). If the logic we uncovered for *C. elegans* applies to the SOD1 G93A model, an elongation inhibitor given at an intermediate concentration to lower protein synthesis by ~30% should rescue some of the disease phenotypes.

Proposing translation inhibitors as potential therapeutics first seems counterintuitive and dangerous. However, besides rapamycin, emetine and homoharringtonine are two additional FDA–approved elongation inhibitors. Emetine is a eukaryote–specific translation inhibitor with a long history of being used in humans, but its nausea-inducing properties make chronic use unsustainable. Homoharringtonine is an elongation inhibitor that only binds vacant ribosomes that can no longer access its binding site once translation has commenced. Essentially it functions as an initiation inhibitor that binds within the ribosomal peptidyl transferase center (Dmitriev et al., 2020). Finally, the tetracyclines, minocycline, and doxycycline, also act as translation inhibitors for both mitochondrial translation and cytoplasmic translation (Molenaars et al., 2020; Mortison et al., 2018; Solis et al., 2018). These examples show that, as long as cytoplasmic translation is reduced rather than wholly blocked, translation inhibitors are tolerable in humans. Thus, translation inhibitors may be suitable for therapeutic development, provided the mode of translational inhibition matches the underlying challenges to proteostasis caused by the disease.

## Supplemental Figure

**S1.**
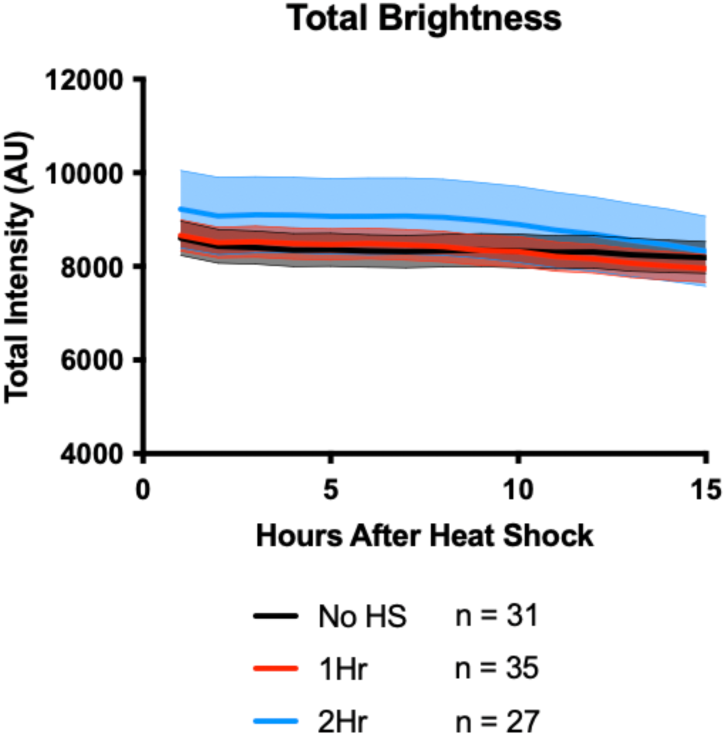
Total fluorescent YFP intensity in PolyQ transgenic animals does not change significantly within the 15 hours imaging of the animals after the HS, showing that the aggregate formation is a redistribution of soluble PolyQ::YFP into aggregation foci. Lines indicate mean, and shading indicates 95% CI.

**S2.**
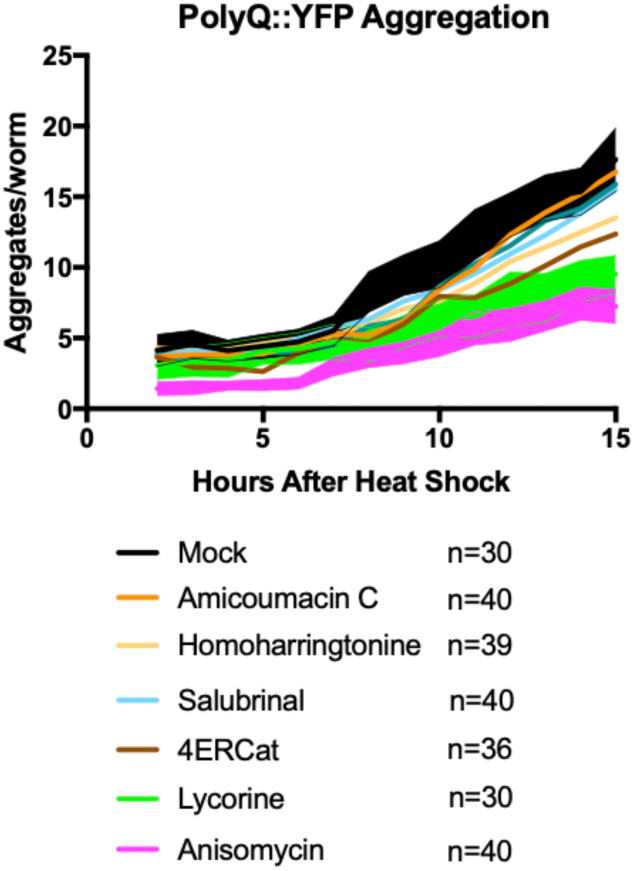
Transgenic AM140 worms treated with indicated chemicals for 12 hours at the late L4 stage, then subjected to 36 °C for 2 hours on day 1 of adulthood. Only anisomycin and lycorine had suppressed aggregate formation as measured in our live imaging protocol. Lines indicate mean, and shading indicates 95% CI for DMSO, anisomycin, and lycorine.

**S3.**
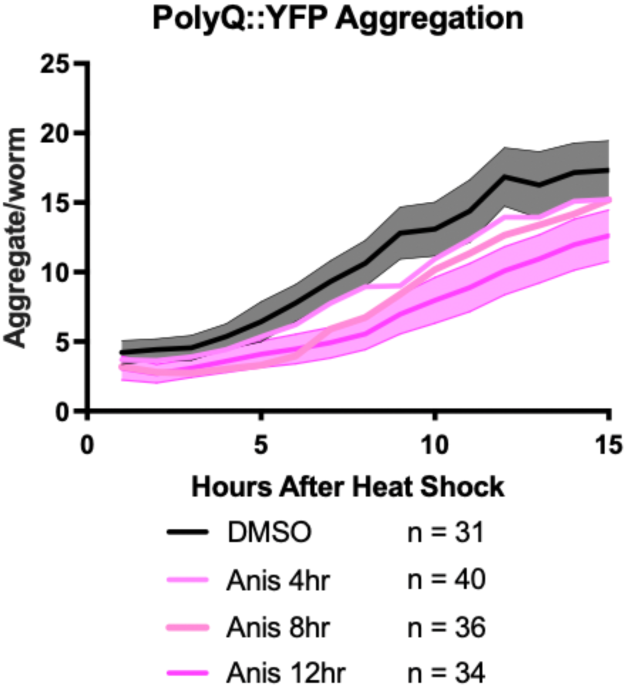
12-hour pre-treatment of anisomycin in PolyQ *C. elegans* was necessary to inhibit aggregation significantly. Lines indicate means, while DMSO and 12-hour treatment with anisomycin (100 μM) include 95% CI as indicated by shading.

**S4.**
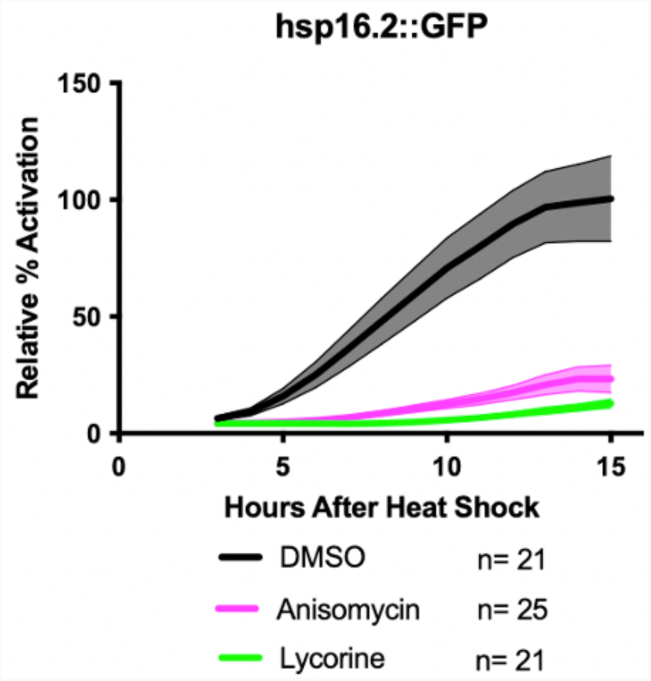
Elongation inhibitors suppressed the HSR as measured by *hsp*-16.2::GFP fluorescence assay. After incubation with the inhibitors for 4 days followed by a 1 hour HS at 36 °C, little to no increase in GFP expression was observed, indicating that all inhibitors block HSR activation at the tested concentration (100 μM). Lines indicate mean, and shading indicates 95% CI.

**S5.**
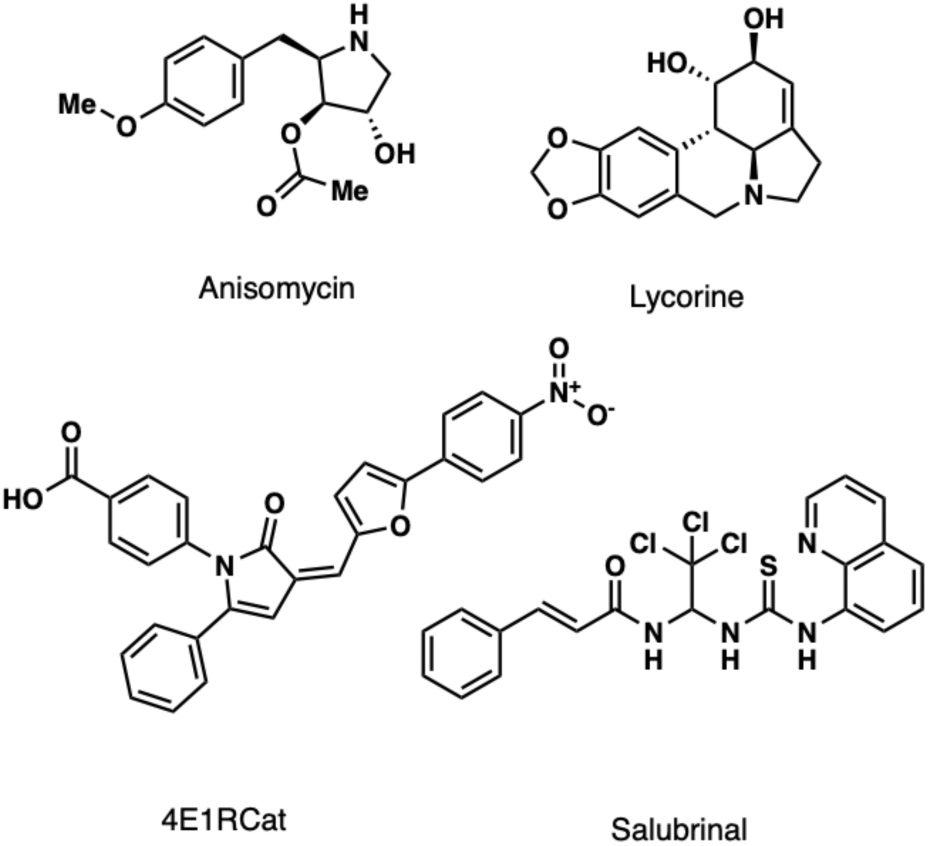
Chemical structures of anisomycin, lycorine, 4E1RCat, and salubrinal.

**Supplemental Table 1:**
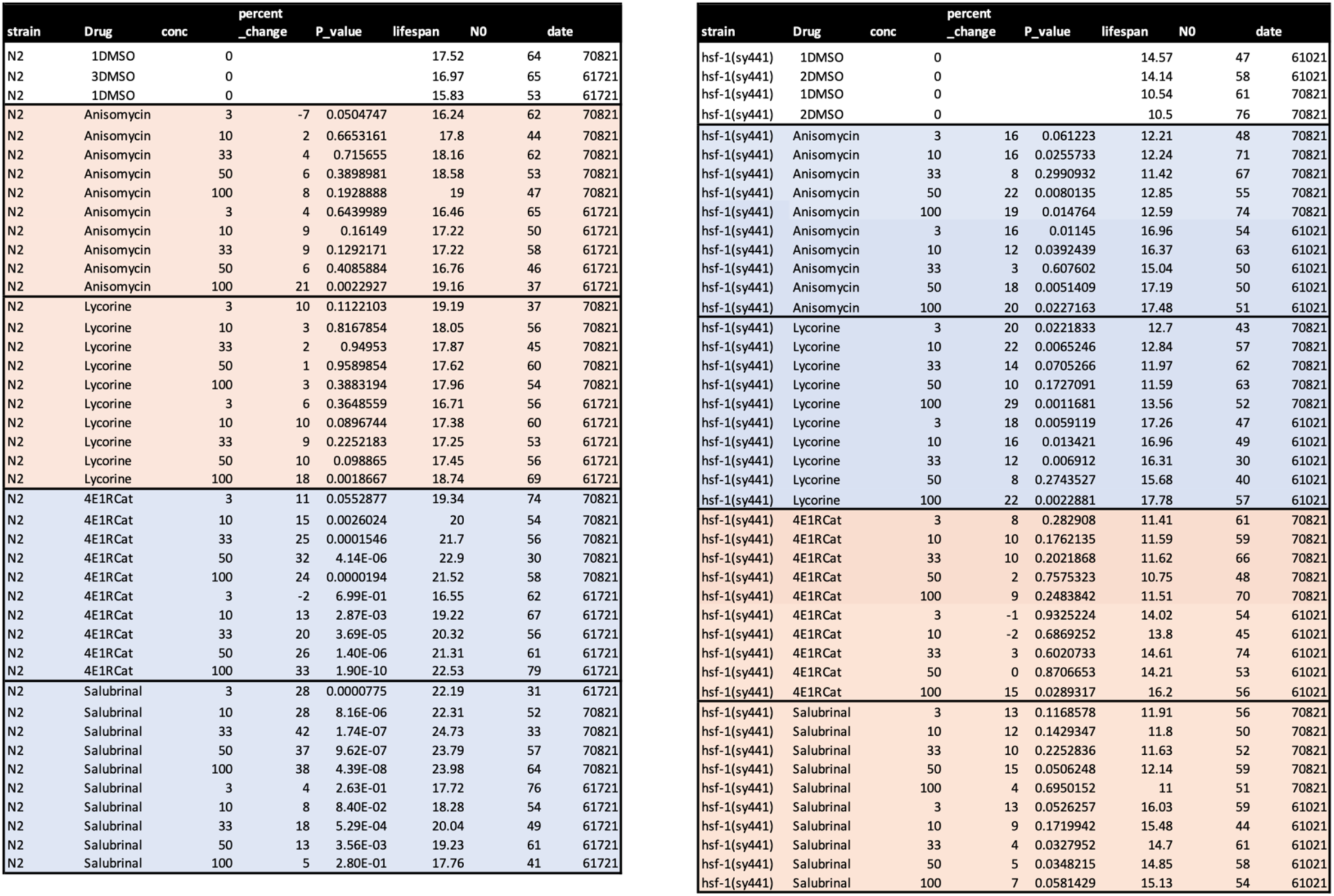
Representative dose-response curves of DMSO, anisomycin, lycorine, 4E1RCat, and salubrinal in 96-well plate lifespan assays, repeated in two complete individual replicates. For N2, only 4ER1RCat and salubrinal robustly increase lifespan (blue), while anisomycin and lycorine either are not effective or slightly toxic (red). For *hsf-1(sy441)*, only anisomycin and lycorine robustly increase lifespan (blue), while 4E1RCat and salubrinal either are not effective or slightly toxic (red).

## Acknowledgments

We would like to acknowledge Drs Anabel Perez–Gomez, Sarah Ly, Jin Lee, Caroline Kumsta, and Malene Hansen for input into the manuscript; Alan To for technical assistance; the Yusupov lab for generating the structures of anisomycin, lycorine, and homoharringtonine in complex with the eukaryotic ribosome; and Dr. Shigefumi Kuwahara (Tohoku University) for providing Amicoumacin C. Grants supported this work to M.P. from the NIH (DP2 OD008398, R21NS107951, R01AG067331), and the Glenn Foundation. K.C. was funded by the Dorris Neuroscience Scholar Fellowship. Some strains were provided by the CGC, which is funded by the NIH Office of Research Infrastructure Programs (P40 OD010440).

## Competing interests

M.P. and K.C are scientific founders and advisors to Cyclone Therapeutics, Inc., a biotech company developing therapeutics targeting translation.

### Lead Contact and Materials Availability

Michael Petrascheck is the Lead Contact and may be contacted at. This study did not generate new unique reagents; however, the natural product Amicoumacin C was obtained as a gift from Dr. Shigefumi Kuwahara, Ph.D. (Tohoku University). This reagent is not available without total chemical synthesis.

### Experimental Model and Subject Details

#### *C. elegans* Strains

The Bristol strain (N2) was used as the wild-type strain. The following worm strains used in this study were obtained from the Caenorhabditis Genetics Center (CGC; Minneapolis, MN): CL2070 [dvIs70 [*hsp*-16.2p::GFP + *rol-6(su1006)*]], AM140 [rmIs132[P*unc-54*::q35::yfp]] and PS3551 [*hsf-1(sy441*)].

### Method Details

#### Worm Maintenance

1000 – 2000 age-synchronized animals were plated into 6 cm culture plates with liquid medium (S-complete medium with 50 mg/mL carbenicillin and 0.1 mg/mL fungizone (Amphotericin B)) containing 6 mg/mL X-ray irradiated Escherichia coli OP50 (1.5 × 10^8^ colony-forming units [cfu]/ml, carbenicillin–resistant to exclude growth of other bacteria), freshly prepared 4 days in advance, as previously described (Solis and Petrascheck, 2011), and were maintained at 20 °C. The final volume in each plate was 7 mL. To prevent self-fertilization, FUDR (5-fluoro-2’-deoxyuridine, 0.12 mM final) (Sigma-Aldrich, cat # 856657) was added 42—45 hours after seeding. At the late L4 stage, either DMSO/drug treatment (100μM unless otherwise stated) was added to each strain.

#### Surface sensing of translation (SUnSET) to analyze the effectiveness of translation inhibitors in C. elegans

Day 1 adult N2 worms were bleached, and eggs were allowed to hatch in S-complete by shaking them overnight. On the next day, 12,000 L1 worms were seeded in a 15 cm plate containing a total volume of 30 mL S-complete with 6 mg/mL OP50 bacteria, 50 μg/mL Carbenicillin, and 0.1 μg/mL Amphotericin B. 6mL of 0.6 mM Fluorodeoxyuridine (FUDR) were added to worms at L4 stage in each plate. 100 μM translation inhibitor was added to worms 2 hours after adding FUDR. After 12 hours, worms were transferred into a 15mL corning tube containing a total volume of 5 mL S-complete with 750 ul 6 mg/mL OP50 bacteria, 0.5 mg/mL puromycin, and 100 μM translation inhibitors. After rotating the corning tubes for 4h, worms were collected into 2 mL cryotubes by washing them with M9 once and with cold PBS 3 times. Worms were flash-frozen in liquid nitrogen and subsequently broken with a beak mill homogenizer (Fisherbrand). Protein concentrations were determined by the Bradford protein assay. 50 mg protein from each sample was loaded for western blot analysis using antibodies against puromycin (Millipore, MABE343) and GAPDH (Proteintech, 10494-1-AP). Antibodies were diluted 1:5,000 in 5% non-fat milk in TBST.

#### Thermotolerance

Age-synchronized N2 or PS3551 [*hsf-1(sy441*)] animals were prepared as above in 6 cm culture plates and treated with water or 100 mM lycorine/anisomycin on day 1. On day 4, 25 – 35 animals were transferred to 6 cm NGM plates in triplicate for each condition and were transferred to the non–permissive temperature of 36 °C. Every hour, survival was scored by lightly touching animals with a worm pick and scoring for movement.

#### *C. elegans* insoluble protein extraction

10,000 N2 worms were sorted into a 15cm liquid culture dish using the COPAS Biosorter. For heat shock-induced aggregation experiments, worms were treated with either DMSO or anisomycin (100 μM) for 12 hours on Day 1 of adulthood, then subjected to a 2-hour heat shock at 36 °C. After 12 hours of recovery, the animals were washed 3 times with S-complete buffer, once with PBS, and then flash-frozen in liquid nitrogen. 500uL of cold lysis buffer (20mM Tris base, 100mM NaCl, 1mM MgCl_2_, pH = 7.4, with protease inhibitors (Roche, 11836153001) was added, and animals homogenized mechanically. An aliquot of this total lysate was saved. In an ultracentrifuge tube, 2 volumes of SDS Extraction buffer (20mM Tris base, 100mM NaCl, 1mM MgCl_2_, pH = 7.4, with protease inhibitors, and 1% SDS) was added to 1 volume of total lysate and was centrifuged at 20,000 x g for 30 minutes. The extraction was repeated 2 times to remove all SDS soluble proteins. The remaining insoluble pellet was suspended in 20uL urea buffer (8M urea, 50mM DTT, 2% SDS, 20mM Tris base, pH = 7.4) and sonicated briefly. 18uL of the insoluble suspension was added to 6uL 4x Laemmli buffer (Biorad, #161-0747) supplemented with 10% 2-mercaptoethanol (Sigma, 60-24-2) and boiled for 5 minutes, then directly loaded onto SDS-Page gel (Biorad, 4569033). Gels were stained with Sypro Ruby according to the manufactures directions and quantified in ImageJ.

For age-associated protein aggregation experiments, the above was repeated with the following changes: Day 1 worms were treated with 100 μM of each compound and allowed to age in liquid culture until Day 8 of adulthood. Following lysis, worms were washed several times with PBS to remove soluble protein.

#### Proteasome dysfunction assay—Survival

Animals were prepared as above in 96 well plates. At the late L4 stage, animals were pretreated with DMSO or 100 μM anisomycin. After 12 hours, the animals were treated with 75 μM bortezomib. On day 8 of adulthood, the percent of animals alive was determined by movement in liquid culture.

#### Proteasome dysfunction assay—Worm Length

Animals were prepared as above in 6cm liquid culture dishes. At the late L4 stage, animals were pretreated with DMSO or 100 μM anisomycin. After 12 hours, the animals were treated with 75 μM bortezomib. Body length was measured on day 3 for *hsf-1(sy441)* or day 5 for PolyQ using the 10x objective with the ImageXpress Micro XL and Metaexpress microscopy software.

#### Heat Shock Induced Aggregation and Stress Response

CL2070 [dvIs70 [*hsp*-*16*.*2p*::GFP + *rol-6*(su1006)]] or AM140 [rmIs132[Punc-54::q35::yfp]] age–synchronized animals were treated with DMSO or 100 μM drug at late L4. 4–12 hours later, on day 1 of adulthood, 1.5 mL of the treated animals were transferred from liquid culture into an Eppendorf tube, washed twice with S-complete, pelleted, then transferred to 6 cm NGM plates using S-complete. Once the animals were completely dry on the NGM plate, they were transferred to a 36 °C incubator, plates upside down, for 1–2 hours.

#### Hydrogel Mounting

Animals were washed from NGM plates using 0.2% HHPPA (2-hydroxy-4’-(2-hydroxyethoxy)-2-methylpropiophenone) (CAS[106797-53-9]) dissolved in S-complete, into a 2 mL Eppendorf tube. Animals were washed twice with 0.2% HHPPA, then suspended in 0.3 mL 0.2% HHPPA. 2.5 uL of this solution was seeded into a single well of a 384–well plate containing 2.5 uL 30% PEG-DA (polyethylene glycol diacrylate, MW=4000, Polysciences, cat # 15246-1) in S-complete. After 5 minutes to allow the solutions to diffuse together, the animals were immobilized by subjecting the 384–well plate UV light using a routine laboratory gel viewer (UVP Dual-Intensity Ultraviolet Transilluminator, high intensity) for 30 seconds. 45 uL S-complete buffer was added on top to prevent desiccation. In general, each well contained 5–10 worms.

#### Imaging and Analysis

Time-lapse brightfield and fluorescence images were taken with a 10x objective using the ImageXpress Micro XL over 15 hours. The number of PolyQ aggregates, or total YFP fluorescence, in the whole worm was determined by analyzing images using a custom pipeline created in CellProfiler.

#### Lifespan Assay

Age–synchronized *C. elegans* were prepared in liquid medium, as described above, and seeded into flat-bottom, optically clear 96–well plates (Corning, 351172) containing 150uL total volume per well, as previously described (Clay and Petrascheck, 2020). Plates contained ~10 animals per well in 6 mg/mL γ–irradiated OP50. Age– synchronized animals were seeded as L1 larvae and grown at 20 °C. Plates were covered with sealers to prevent evaporation. To prevent self-fertilization, FUDR (0.12mM final) was added 42—45 hours after seeding. Drugs were added on day 1 of adulthood. When used, DMSO was kept to a final concentration of 0.33% v/v.

### Quantification and Statistical Analysis

#### Aggregation and Induction of the HSR

The number of n represents the total number of animals over three individual experiments. For the paired– time-lapse data generated, we chose to depict the 95% confidence interval, which was calculated using Graphpad Prism, to show differences in treatment.

#### Percent Survival — Thermotolerance assay

The number of n represents the total number of animals over the 3 individual replicates shown as the average percentage survival and SEM, calculated using Graphpad Prism. Significance was determined by using a row– matched two–way ANOVA with Šídák multiple comparisons test.

#### Worm Length — Proteasome dysfunction assay

The number of n represents the total number of animals whose length was measured in one experiment. Depicted are the mean and standard deviation calculated using Graphpad Prism. Significance was determined by the two-tail unpaired t-test. Similar results were observed across 3 independent experiments.

#### Lifespan Assay

Survival was scored manually by visually monitoring worm movement using an inverted microscope 3 times per week. Statistical analysis was performed using the Mantel–Haenzel version of the log-rank test as outlined in Petrascheck and Miller (Petrascheck and Miller, 2017).

## Data and Code Availability

The software used in this study (Cell Profiler) is available at cellprofiler.org

